# Combined effects of matrix stiffness and obesity-associated signaling directs progressive phenotype in PANC-1 pancreatic cancer cells *in vitro*

**DOI:** 10.1101/2024.06.11.598541

**Authors:** A.E. Jones, J.F. Netto, T.L. Foote, B.N.K. Ruliffson, C.F. Whittington

## Abstract

Obesity is a leading risk factor of pancreatic ductal adenocarcinoma (PDAC) that contributes to poor disease prognosis and outcomes. Retrospective studies have identified this link, but interactions surrounding obesity and PDAC are still unclear. Research has shifted to contributions of fibrosis (desmoplasia) on malignancy, which involves increased deposition of collagens and other extracellular matrix (ECM) molecules and increased ECM crosslinking, all of which contribute to increased tissue stiffening. However, fibrotic stiffening is underrepresented as a model feature in current PDAC models. Fibrosis is shared between PDAC and obesity, and can be leveraged for *in vitro* model design, as current animal obesity models of PDAC are limited in their ability to isolate individual components of fibrosis to study cell behavior. In the current study, methacrylated type I collagen (PhotoCol®) was photo-crosslinked to pathological stiffness levels to recapitulate fibrotic ECM stiffening. PANC-1 cells were encapsulated within PhotoCol®, and the tumor-tissue constructs were prepared to represent normal (healthy) (∼600 Pa) and pathological (∼2000 Pa) tissues. Separately, human mesenchymal stem cells were differentiated into adipocytes representing lean (2D differentiation) and obese fat tissue (3D collagen matrix differentiation), and conditioned media was applied to PANC-1 tumor-tissue constructs. Conditioned media from obese adipocytes showed increased vimentin expression, a hallmark of invasiveness and progression, that was not seen after exposure to media from lean adipocytes or control media. Characterization of the obese adipocyte secretome suggested that some PANC-1 differences may arise from increased interleukin-8 and -10 compared to lean adipocytes. Additionally, high matrix stiffness associated induced an amoeboid morphology in PANC-1 cells that was not present at low stiffness. Amoeboid morphology is an accessory to epithelial-to-mesenchymal transition and is used to navigate complex ECM environments. This plasticity has greater implications for treatment efficacy of metastatic cancers. Overall, this work 1) highlights the importance of investigating PDAC-obesity interactions to study the effects on disease progression and persistence, 2) establishes PhotoCol® as a matrix material that can be leveraged to study amoeboid morphology and invasion in PDAC, and 3) emphasizes the importance of integrating both biophysical and biochemical interactions associated within both pathologies for *in vitro* PDAC models.

## 1. INTRODUCTION

Pancreatic ductal adenocarcinoma (PDAC) is set to become the second deadliest cancer by 2030.^1^ This ascension from third deadliest to second likely results from factors such as the lack of effective screening tools and targeted PDAC therapies beyond erlotinib, combined with increased treatment and screening efficacy for other cancers. Moreover, there are several unmodifiable (e.g., family history, ethnicity, age) and modifiable (e.g., obesity, diabetes, tobacco use) risk factors associated with PDAC, some of which are on the rise.^2–4^ Obesity is one of the primary modifiable PDAC risk factors, with high body mass index (BMI) individuals having a 30% increased risk for developing pancreatic cancer when compared to lower BMI counterparts.^5^ In addition, fibrosis within PDAC tumors (desmoplasia) is a hallmark of the disease that imparts numerous biochemical and biophysical changes within the tumor microenvironment.^6–9^The adipose microenvironment in obesity undergoes similar biochemical and biophysical alterations to the ECM that have been shown to induce tumorigenesis in certain cancers (e.g., breast, pancreatic).^10,11^ However, there remains a gap in how adipose cell signaling within obesity conditions drive and sustain malignancy after cancers like PDAC develop.

Bidirectional signaling between obese adipose tissue and pancreatic tumor tissue plays a role in progressing both pathologies. *In vitro* studies of indirect co-culture and conditioned media from pancreatic cancer cells have been shown to promote adipocyte dysfunction that is characterized by loss of lipid droplet formation and adoption of a fibroblast-like phenotype.^12,13^ When cancer-associated stromal cells develop a fibroblast-like phenotype, they contribute to the formation of an inflammatory environment characterized by the secretion of inflammatory cytokines (e.g. IL-6, IL-8, IL-1β).^14,15^ Mouse models of PDAC and obesity have also demonstrated that weight-gain creates an inflammatory environment that aids cancer progression .^15–17^ However, *in vivo* models are limited in their ability to isolate individual biophysical and biochemical elements within the microenvironment of either PDAC or obesity that contribute to disease progression, which limits the ability to identify potential targets or specific intervention strategies.

One shared feature of the PDAC and obesity tissue microenvironments is altered stiffness brought on by increased fibroinflammatory activity. PDAC is characterized by a dense extracellular matrix (ECM) that accounts for 90% of the tumor’s mass.^18^ During fibrotic progression of the disease (i.e., desmoplasia) activated stromal populations—primarily cancer-associated fibroblasts (CAFs) and pancreatic stellate cells, (PSCs)—deposit increasing amounts of ECM macromolecules (e.g. type I collagen, hyaluronic acid, etc.) and induce enzymatic crosslinking (e.g., lysyl oxidase) of ECM proteins, which raises the bulk stiffness of the diseased tissue. Normal pancreatic tissue stiffness is ∼ 0.5-1kPa, while PDAC tumor stiffness falls within a range of ∼1-8kPa.^19,20^ Fibrosis also occurs in adipose tissue with obesity. Adipocytes compose the parenchymal fraction of adipose tissue. As adipocytes increase in number and size, increased metabolic demands form a hypoxic environment that creates an inflammatory environment. Unresolved inflammation in turn encourages ECM remodeling.^21^ Increased collagen deposition and lysyl oxidase activity within obese adipose tissue results in an increase in ECM stiffness to ∼2-4 kPa in healthy from ∼2-11 kPa during obesity.^22–26^

In both pathologies—PDAC and obesity—resident cells sense and respond to changes in the surrounding microenvironment. Extracellular matrix stiffness has been found to impact changes in pancreatic cancer cell phenotype and disease progression.^7,27–30^ *In vitro* models of PDAC that have focused on altering ECM stiffness properties have shown that increasing matrix stiffness can promote epithelial-to-mesenchymal transition (EMT) in PDAC cells ^27,29–32^. The gradual loss of epithelial markers (e.g., E-cadherin) that promote cell-cell adhesion and progressive gain of mesenchymal markers (e.g., Vimentin) that increase migratory properties are also accompanied by morphological changes as cells transition from rounded epithelial cells to elongated mesenchymal cells. ^33,34^ Collectively, these phenotypic changes can be predictive of progressive behaviors such as extracellular migration and invasion in solid cancers.^33,35^ *In vitro* studies have shown that introducing microenvironmental changes associated with fibrosis, such as increasing matrix density and stiffness, leads to adipocyte dysregulation. ^36–39^This dysregulation is associated with a fibroblast-like phenotype characterized by alpha smooth muscle actin (α-SMA) upregulation and loss of lipid droplet formation. Therefore, shared microenvironmental changes (i.e., ECM stiffening) within both pathologies—PDAC and obesity— can be leveraged to better understand the role ECM stiffness plays in changing cellular responses and paracrine signaling in the PDAC tumor microenvironment.

In this study we investigated the impact of ECM stiffness on altering the response to obesity-associated signaling in PDAC as it relates to disease progression and persistence. To evaluate the influence of these interactions, we used 1) a methacrylated type I collagen hydrogel capable of photo-crosslinking (PhotoCol®, Advanced BioMatrix, Inc., Carlsbad, CA) to expose PANC-1—a mesenchymal PDAC cell line—to increasing matrix stiffness associated with fibrosis and pancreatic cancer progression and 2) conditioned media from lean or obese adipocytes to expose PDAC cells to adipocyte signaling. We hypothesized that increasing matrix stiffness and exposure to obese adipocyte conditioned media would increase cell malignant phenotype related to irregular cell morphologies and vimentin expression. After generating obese adipocytes with 3D differentiation within a high concentration (6 mg/mL) collagen hydrogel rather than 2D differentiation that yields lean adipocytes, we applied conditioned media from obese and lean adipocytes to PANC-1 cells encapsulated within low (∼600 Pa) and high (∼2 kPa) stiffness PhotoCol® matrices. Results showed that PANC-1 cells show more signs of malignancy via increased vimentin expression when exposed to obesity conditioned media, independent of matrix stiffness, but did not exhibit differences in cell shape metrics associated with elongation and cell spread. We also did not see morphological signs of malignancy or epithelial-to-mesenchymal transition with increased matrix stiffness, which was unexpected. However, we observed PANC-1 cells in stiffer matrices with a more amoeboid-like morphology, suggesting that cells were undergoing a mesenchymal to amoeboid transition as malignancy persists. We concluded that our study results reveal a role for obesity signaling in PDAC that does not depend on matrix stiffness; however, our use of PhotoCol® creates an densified matrix environment that offers an opportunity to study an alternate mechanism of disease progression through amoeboid invasion.

## 2 RESULTS

### 2.1 PANC-1 cells in 2D undergo morphological changes with adipocyte conditioned media exposure

In advance of performing conditioned media exposure studies in 3D, we evaluated the impact of adipocyte signaling on PANC-1 cells in 2D. To determine the appropriate concentration of adipocyte conditioned media (ACM) for exposure studies, PANC-1 cells were cultured in different volumetric ratios (50% and 75% v/v) of ACM to PANC-1 growth media and assessed for morphological changes with F-actin staining (Figure 1A). Regardless of media condition, cells maintained excellent viability (>90%) (data not shown), and we proceeded with 75% v/v ACM (ACM-75) for subsequent studies since it contained a higher proportion of adipocyte signaling content. Quantitative cell analysis performed in CellProfiler ^TM^ was used to extract the following shape metrics from individual PANC-1 cells: Compactness, Eccentricity, Extent, Form factor, and Solidity, based on their potential to predict cell invasion.^33^ This analysis was also performed to align with shape quantification for subsequent 3D studies. Analysis revealed that ACM-75 exposure resulted in PANC-1 cells with a significantly lower average form factor compared to control cells, which is indicative of more cell spread and cell protrusion with exposure to adipocyte signaling. These cells also had a significantly higher eccentricity, which means that cells were more elongated. (Figure 1B). None of the other shape parameters—extent, compactness, solidity—showed significance, which was unexpected. However, both an increase in cellular protrusion and cell elongation are associated with a mesenchymal phenotype that aids cell migration and invasion *in vitro*^33,34,40^ , which suggests that adipocyte signaling may be important for sustaining or promoting a more invasive or progressive cell phenotype.

**Figure 1:**
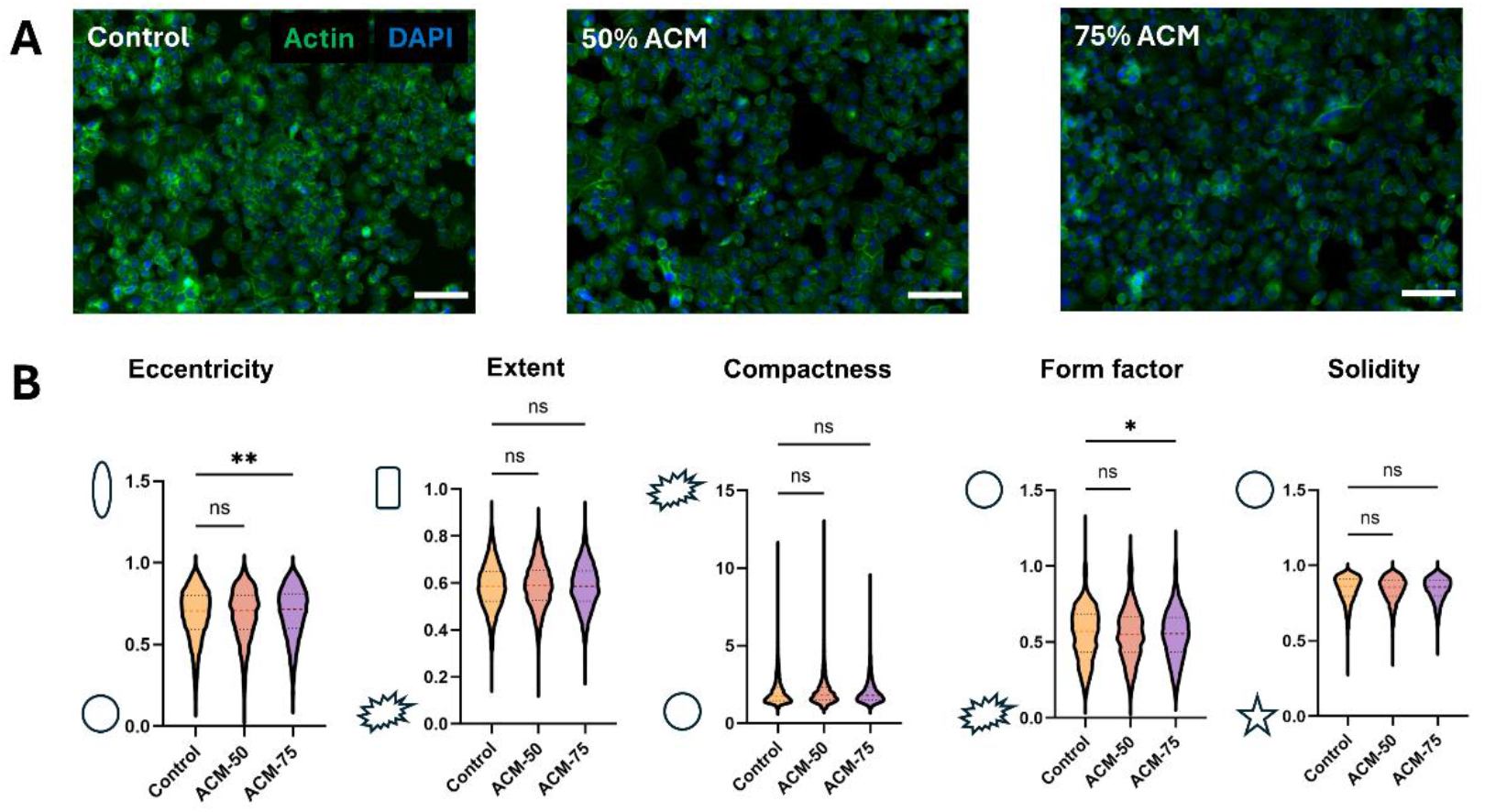
2D PANC-1 morphology with adipocyte conditioned media (ACM) exposure. A) Representative fluorescence images of PANC-1 cells treated with control media, ACM-50, and ACM-75 after 48 hours (green = F-actin, blue = nucleus; Scale bar = 100 µm). B) Quantification of cell shape parameters related to cell elongation (eccentricity, extent) and cell spread (compactness, form factor, solidity) for PANC-1 cells in control media (n=6148), ACM-50 (n=6222), and ACM-75 (n=5042). Data are represented with violin plots to show distribution and variability. Groups were compared with a one-way ANOVA with Dunnett’s multiple comparisons (p ≤ 0.05 (*), p ≤ 0.01 (**), p ≤ 0.001 (***), p < 0.0001 (****), ns = no significance).

### 2.2. Adipocytes secrete obesity-associated factors when cultured within 3D type I collagen (high concentration)

Adipocytes differentiated *in vitro* from mesenchymal stem cells on 2D without additional additives beyond adipogenic factors are meant to be representative of healthy functioning adipocytes (i.e., lean fat) with the associated characteristic markers (e.g., lipid droplet formation, adiponectin secretion, low leptin secretion). To study interactions between PDAC and obesity, we needed adipocytes that are more associated with an obese phenotype. To generate these cells, we differentiated adipocytes in a high concentration type I collagen hydrogel (6 mg/ml; non-methacrylated telo-collagen) to mimic increased collagen deposition that occurs during obesity.^21,36,41^ We verified adipokine expression within conditioned media samples qualitatively (Figure 2A, B) and quantitatively (Figure 2C). Results from a human adipokine array blot verified that conditioned media from adipocytes differentiated in 2D (ACM) contained adiponectin, which is associated with lean fat (Figure 2A). Obesity conditioned media (OCM) from adipocytes differentiated within collagen hydrogels maintained similar levels of adiponectin while also showing increasing amounts of key obesity biomarkers. Selected targets represented the highest fold change (> 2-fold change) between lean and obese adipocytes secretion, except for IL-6 that was included based on association with obesity. Samples of OCM contained significantly higher levels of resistin, c-reactive protein (CRP), and epithelial neutrophil activating peptide (ENA-78/CXCL5) (Figure 2C). Both resistin and ENA-78 confer insulin resistance to adipocytes and increase inflammatory cytokine secretion (Il-8, IL-6, TNF-α) from white adipose tissue *in vitro*.^42–44^ Increased resistin expression has also been observed in poorly differentiated PDAC tumors and correlated to poor outcomes in PDAC. Increased CRP production results from an increase in body mass index (BMI) and similarly tailors the inflammatory environment of obesity.^45,46^ Additionally, we observed significant increases in interleukins 8 and 10 (IL-8, IL-10), which are pro-inflammatory and anti-inflammatory cytokines, respectively, that are both upregulated in obesity.^47–49^ We expected to see increased levels of IL-6, since it has traditionally been associated with obesity^48^ and increased levels of resistin and ENA-78. However, IL-6 levels in our OCM samples were variable, resulting in no significant change in average protein secretion between lean and obese adipocytes. Though, there is evidence to suggest that serum IL-6 levels can be attenuated by increasing IL-10 secretion in obese patients^50^, which may explain our observations.

**Figure 2:**
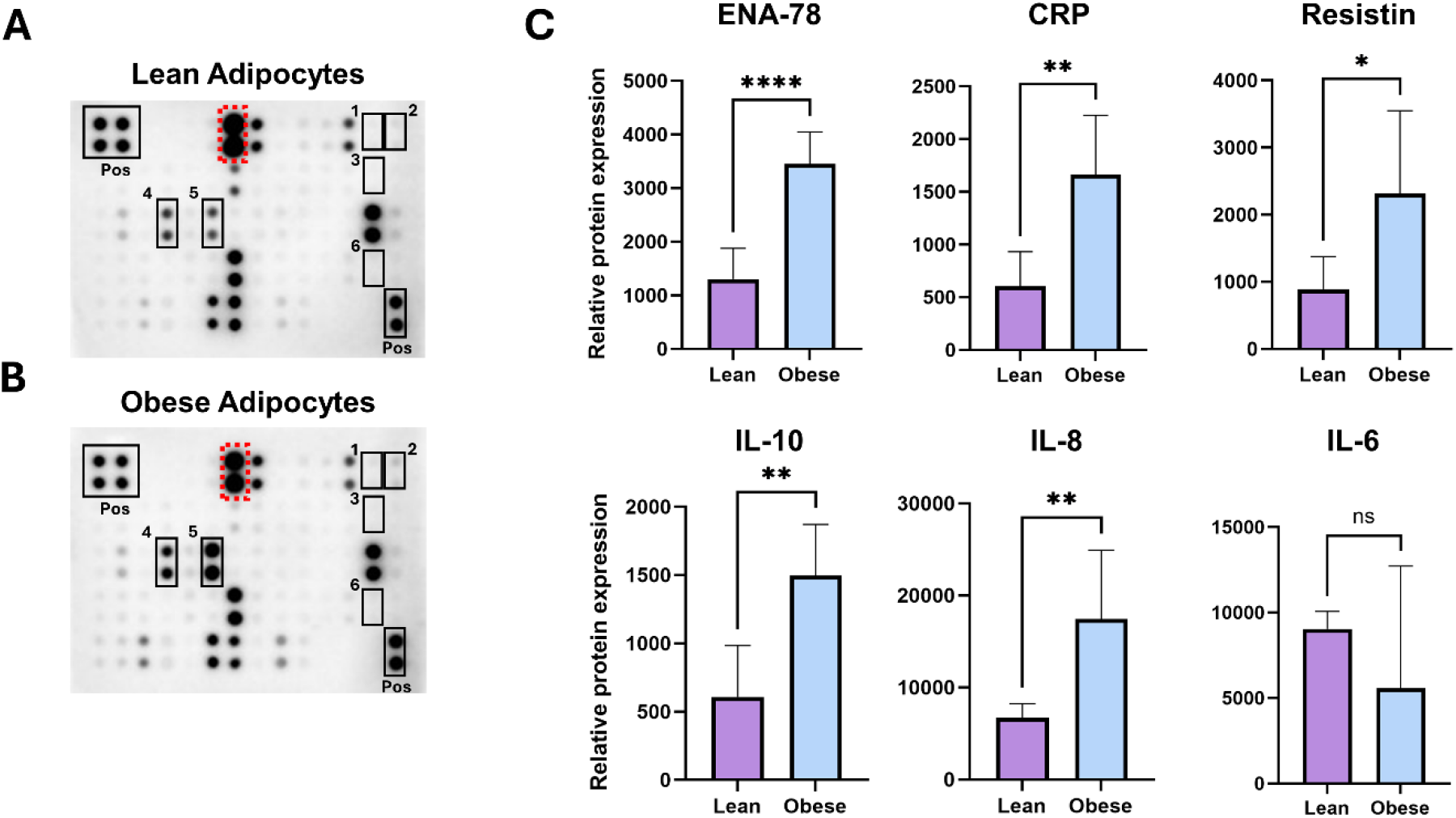
Conditioned media characterization from adipocytes with lean and obese phenotypes. Images of human adipokines array blots used to characterize conditioned media from A) lean and B) obese adipocytes. Red dotted box shows location of adiponectin. Black boxes show locations for obesity markers: 1 – c-reactive protein (CRP), 2 – epithelial neutrophil-activating protein 78 (ENA-78), and 6 – resistin and inflammatory/anti-inflammatory cytokines: 3 – interleukin 10 (IL-10), 4 – interleukin 6 (IL-6), and 5 – interleukin 8 (IL-8). Boxes labeled with “Pos” are the positive array controls. C) Quantified relative protein expression measured by densiometric analysis of adipokine markers for normal and obese adipocyte conditioned media. Bars represent mean+SD. Significance was measured with a student’s t-test (N=3; p ≤ 0.05 (*), p ≤ 0.01 (**), p ≤ 0.001 (***), p ≤ 0.0001 (****), ns = no significance).

### 2.3 Changes in vimentin expression are associated with obesity signaling, not matrix stiffness or cell shape

*In vivo*, primary PDAC tumors are surrounded by a dense and fibrillar ECM. To recapitulate these changes *in vitro*, PANC-1 cells were encapsulated within PhotoCol®—a methacrylated type I collagen hydrogel—with photo-crosslinking (high stiffness, ∼2 kPa) or without photo-crosslinking (low stiffness, ∼600 Pa) and exposed to ACM-75, OCM-75, or control media. Samples were then stained for qualitative (Figure 3, Figure 4) and quantitative (Figure 5) assessment of morphology (via F-actin) and vimentin expression. At low stiffness, PANC-1 cells grew in both multicellular clusters and as individual cells regardless of media exposure conditions (Figure 3). Culture with ACM-75 produced no significant changes in cell morphology compared to control media. However, exposure to OCM-75 produced some PANC-1 cells with long protrusions. Cells grown in high stiffness matrices displayed altered F-actin organization (Figure 4), that manifests as more punctate actin staining when compared to low stiffness culture. For low and high stiffness samples, vimentin staining appeared more prominent in samples exposed to OCM compared to ACM or control media. Additionally, some cells had cortical organization of vimentin in high stiffness control samples, but treatment with OCM resulted in uniform vimentin expression throughout the cell regardless of sample stiffness..

**Figure 3:**
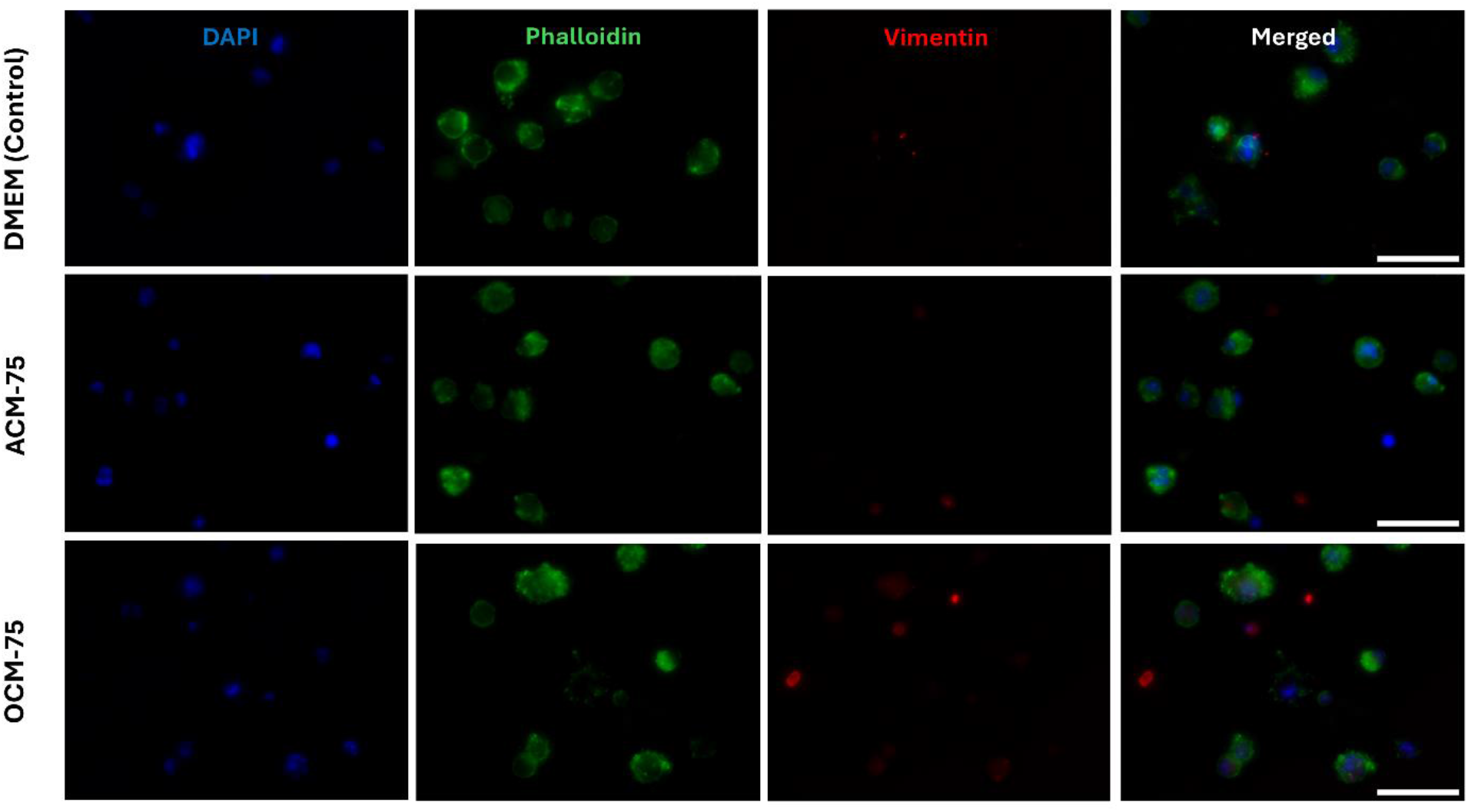
3D PANC-1 morphology in low stiffness methacrylated type I collagen hydrogels. Representative fluorescence images of PANC-1 cells encapsulated in uncrosslinked PhotoCol® hydrogels (low stiffness) and exposed to control PANC-1 media, ACM-75, or OCM-75. Samples were stained to assess morphology (green = F-actin) and expression of mesenchymal marker (red = vimentin). All samples were counterstained to visualize the nucleus (blue = DAPI). Images represent 2D maximum projection of z-stacks (20X objective; 100-200 µm depth, 7 µm pitch; Scale bar = 100 µm).

**Figure 4:**
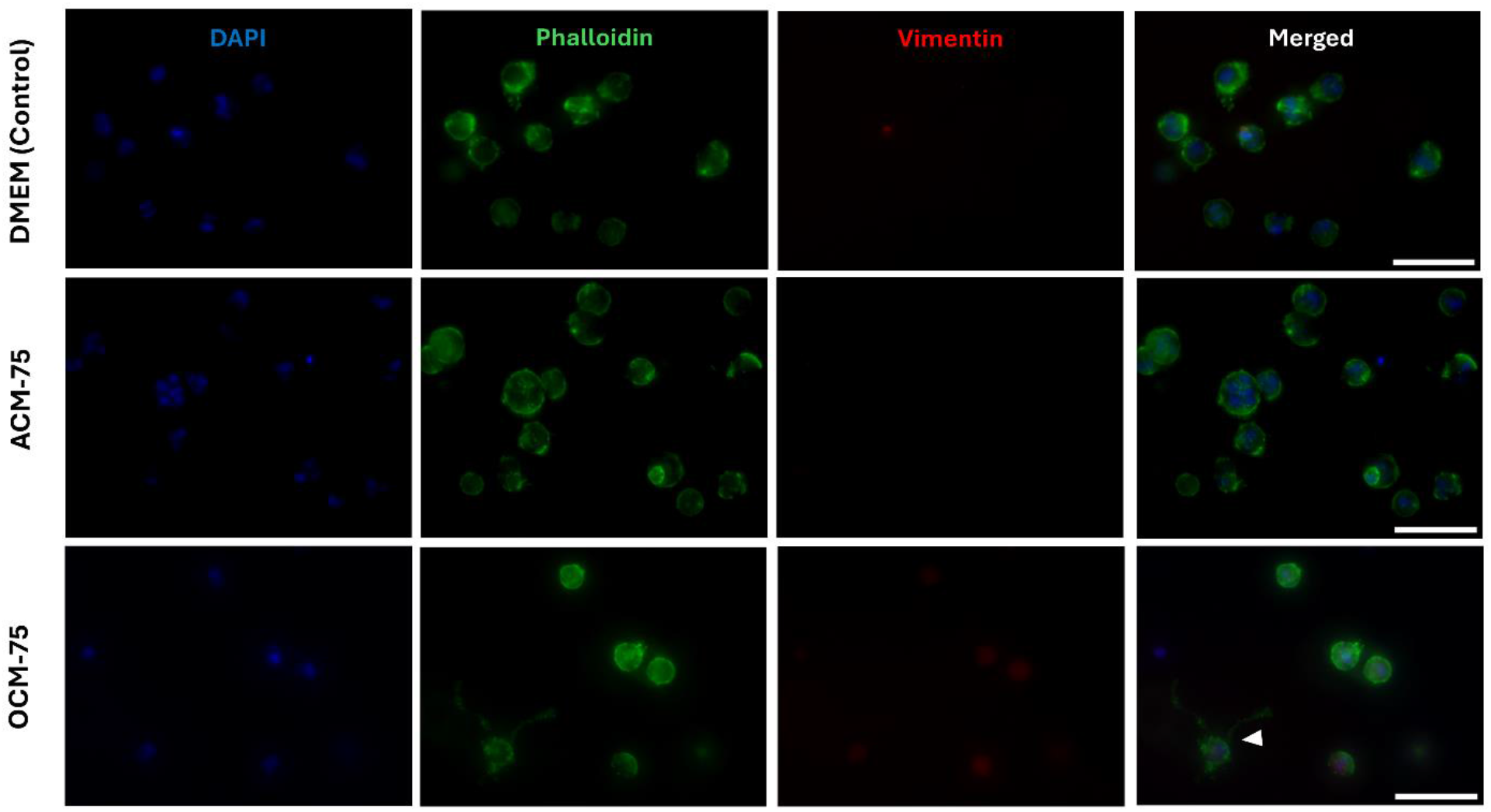
3D PANC-1 morphology in high stiffness methacrylated type I collagen hydrogels. Representative fluorescence images of PANC-1 cells encapsulated in photo-crosslinked PhotoCol® hydrogels (high stiffness) and exposed to control PANC-1 media, ACM-75, or OCM-75. Samples were stained to assess morphology (green = F-actin) and expression of mesenchymal marker (red = vimentin). All samples were counterstained to visualize the nucleus (blue = DAPI). Images represent 2D maximum projection of z-stacks (20X objective; 100-200 µm depth, 7 µm pitch; Scale bar = 100 µm).

**Figure 5:**
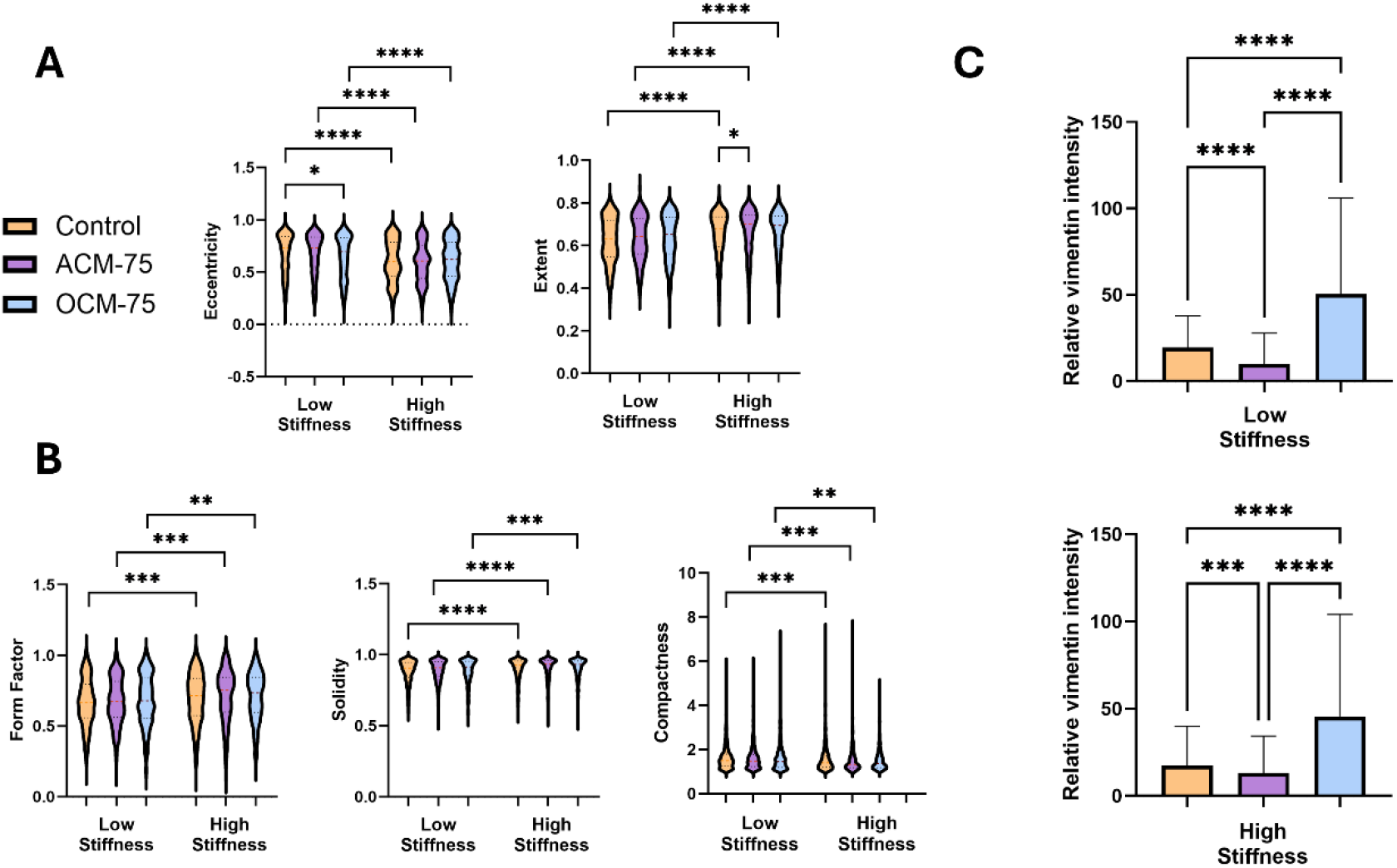
Quantification of cell shape metrics and vimentin expression. A) Measures of PANC-1 elongation (eccentricity, extent) and B) cell spread (form factor, solidity, compactness) for control media (n=113; n=596), ACM-75 (n=1139; n=381), and OCM-75 (n=1182; n=721) at a low or high matrix stiffness, respectively. C) Relative vimentin intensity, as measured by integrated intensity values from fluorescently stained samples, was quantified. Bars represent mean + SD. Significance was measured with a Kruskal-Wallis ANOVA with Dunn’s multiple comparisons within stiffness levels and a Mann-Whitney U for comparison between stiffness levels (p < 0.05 (*), p ≤ 0.01 (**), p < 0.001 (***), p ≤ 0.0001 (****)).

Quantitative analysis of the five shape metrics of cell elongation and spread, as performed for Figure 1, were evaluated. At low matrix stiffness, there was a significant difference in cell eccentricity and form factor between control media and OCM-75 exposed cells (Figure 5A-B). Interestingly, these changes are associated with more uniform cell boarders and less elongated cells. This result contradicts our original hypothesis, as we expected OCM to promote more protrusive and elongated cell morphology, as some cells stimulated with obese conditioned media formed long protrusions. Morphological analysis for samples cultured within stiff matrices revealed that similar changes were observed at a high matrix stiffness as with low matrix stiffness, with PANC-1 cells presenting a significantly higher extent, solidity and lower form factor following exposure to either ACM-75 or OCM-75. These results indicate that cells grown under high stiffness conditions exhibited significantly more uniform cell shape than low stiffness counterparts for all media types.

We measured vimentin expression between low and high stiffness matrix conditions and the different conditioned media formulations (Figure 5C). Vimentin expression, as measured by relative vimentin intensity, was not significantly different between high and low stiffness matrices for each media formulation as only slight increased or decreased levels were observed. This result was surprising given that others have observed significant increases in vimentin expression with increased matrix stiffness for pancreatic cancer cell lines.^27^ However, for our study, significant differences in vimentin were observed with different media exposures, independent of stiffness. Compared to control media, vimentin expression decreased significantly in ACM-75 exposed PANC-1 samples and increased significantly in OCM-75 exposed samples. The vimentin expression in samples exposed to OCM-75 was also significantly higher than samples exposed to ACM-75. Collectively, these results suggest that adipocytes with an obesity-associated phenotype contribute to PDAC malignancy and progression via the more traditional EMT hallmark—increased expression of vimentin as an indicator of mesenchymal phenotype. Moreover, we established that our OCM contains significantly higher amounts of cytokines Il-10 and Il-8 than ACM, which could contribute to the increased vimentin expression we observed, as both cytokines promote EMT.^51,52^

As previously described, all shape metrics associated with cell spread—form factor, solidity, compactness—were unaffected by media formulation. For elongation metrics—eccentricity and extent—the only samples that had any significant differences compared to control media were OCM-75 at low stiffness for eccentricity and ACM-75 at high concentration for extent. Eccentricity changes following OCM-75 stimulation resulted in less elongated cells following conditioned media exposure. The same interpretation is true for extent following ACM-75 exposure. A higher extent means less elongated cells, but this measure is more sensitive to cell protrusions. It is likely that both the high density of collagen fibrils in our hydrogel due to high collagen concentration (7.262 mg/mL) and additional crosslinking led to increased physical confinement of PANC-1 cells. This confinement resulted in more rounded cells and blebbing of the actin cytoskeleton that could indicate an amoeboid cell morphology (Figure 6). Physical confinement of cancer cells by a 3D environment can lead to amoeboid migration to navigate complex microstructure or when proteolytic degradation is not an option, .^53,54^ Photo-crosslinking of methacrylated collagen introduces degradation resistant regions that can significantly decrease degradation time *in vitro* compared to uncrosslinked counterparts.^55,56^ An amoeboid cell morphology is part of the larger cell migration/invasion paradigm (Figure 6A), wherein mesenchymal cells can develop an ameboid phenotype.^57–59^ Therefore, we may have created an environment that reflects the amoeboid transition as an alternative form of PDAC progression within a desmoplastic environment. Overall, these results suggest that obese adipocytes do contribute to PDAC malignancy by increasing expression of vimentin—a mesenchymal marker. However, this change in vimentin is not associated with changes in cell shape metrics associated with cell elongation or spread that are more dictated by matrix stiffness and possible confinement.

**Figure 6:**
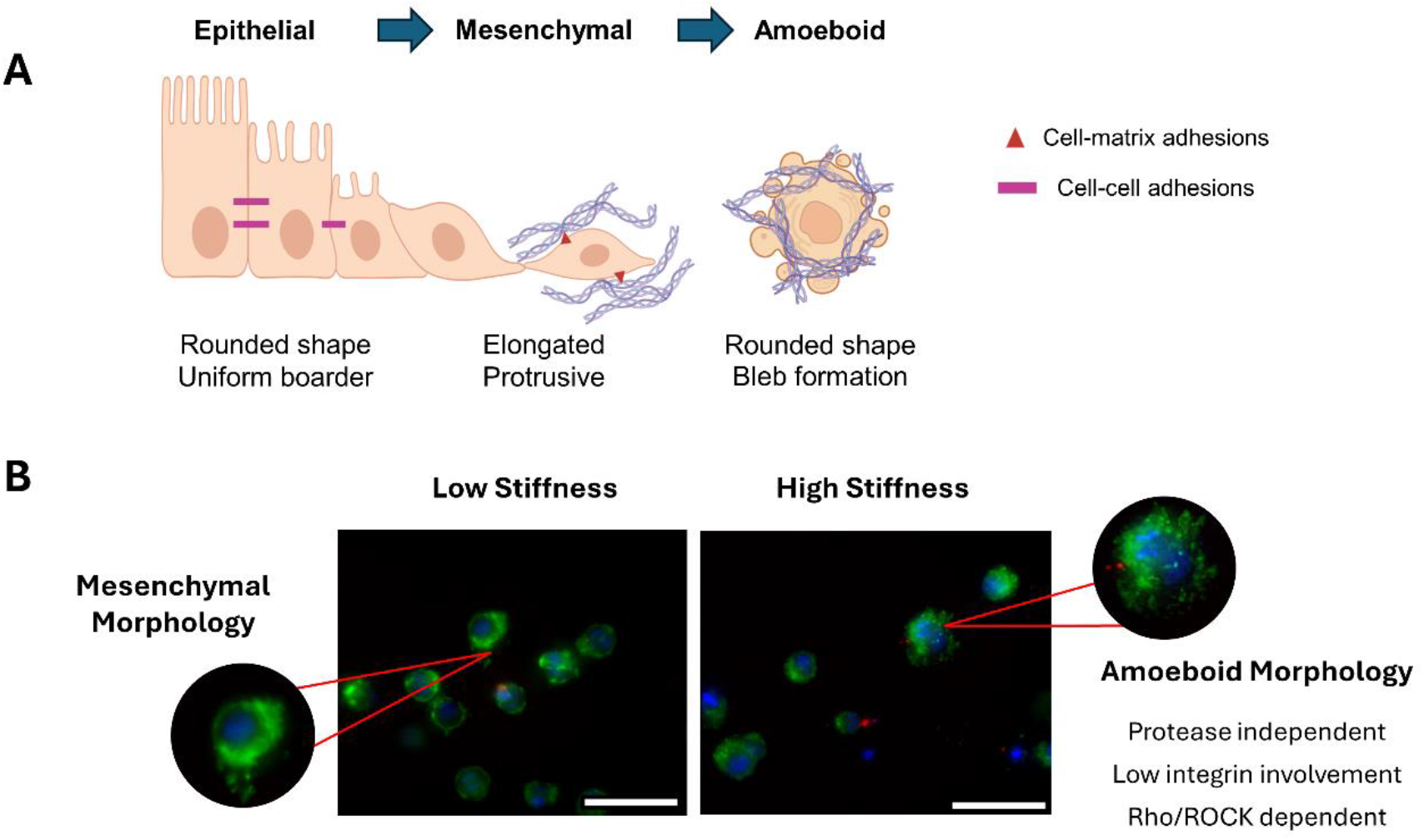
Mesenchymal-to-amoeboid transition in the cell migration cascade. A) Schematic overview of mesenchymal-to-amoeboid transition (MTA) migration as a sequence of traditional epithelial to mesenchymal transition (EMT) (Image created with BioRender). B) PANC-1 cells in a methacrylated type-I collagen hydrogel with normal growth medium under low (left panel) and high (right panel) matrix stiffness with zoomed in portions to show observed differences in mesenchymal and amoeboid morphologies. (green = F-actin, blue = nucleus; Scale bar = 100 µm).

## 3. DISCUSSION

Our study involves two key factors that impact PDAC—desmoplasia and obesity—and we integrated them into an *in vitro* model by culturing PANC-1 cells on methacrylated type I collagen (PhotoCol®) at different stiffness levels and exposing samples to conditioned media from lean or obese adipocytes. We found that adipocytes representative of either lean or obese fat produced different signaling factors based on their phenotype. Unlike lean adipocytes, obese adipocytes increased malignancy of PANC-1 cells independent of matrix stiffness. Additionally, while matrix stiffness did not differentially alter vimentin expression within PANC-1 cells as a marker of EMT and traditional pathway for disease progression, high matrix stiffness appeared to promote amoeboid cell morphologies, which has been identified as an alternative pathway for malignancy.

The malignant influence of the fibrotic PDAC ECM that arises during desmoplasia has prompted the development of matrix targeting therapies. While these therapies showed promising results in animal models, they resulted in poor patient outcomes in clinical trials,^60^ because fibrosis plays a dual role in cancer progression. On the tumor promoting side, the dense stroma can inhibit diffusion of therapeutics and induce more mesenchymal behaviors in cells.^61,62^ On the tumor suppression side, the matrix provides physical confinement that inhibits migration of less invasive cancer cells. When this barrier is removed by matrix ablating therapies, unrestrained tumor cell migration is permitted. Moreover, when we consider the factors that contribute to the densification of the stroma, the primary mechanisms are excess ECM deposition and enzymatic crosslinking of ECM collagens via lysyl oxidase (LOX). LOX expression is upregulated in solid cancers and contributes to matrix stiffness, but the influence of ECM stiffening via crosslinking rather than ECM deposition is understudied as a driver of cancer progression.^30,62,63^ Therefore, it is beneficial to have models that assess the influence of matrix stiffening independent of matrix deposition on cell behavior. We addressed this gap by utilizing a PhotoCol® matrix, to study cell behaviors in a system that decouples collagen concentration and matrix stiffness alongside the influences of obesity signaling with the tumor microenvironment. Additionally, photo-crosslinking was meant to mimic LOX-mediated crosslinking that occurs during fibrosis *in vivo*,^30,62,63^ albeit on a faster timeline than crosslinking occurs *in vivo*.

While a significant increase in mesenchymal phenotype in response to increasing matrix stiffness was not observed in our study, we did see ameboid-like cell morphology in high stiffness matrices that were not present in low stiffness matrices. Traditionally, increased cellular protrusion and elongation is associated with increased malignancy and invasion through EMT.^30^ However, these results can vary depending on the properties of the model system being used, such as dimensionality, cell type, biomaterial formulation, and matrix stiffness.^64,65^ Moreover, epithelial cells are primarily associated with collective migration and invasion, while mesenchymal cells migrate and invade as individual cells. However, those classifications are typically associated with traditional EMT behavior (i.e., loss of epithelial markers and gain of mesenchymal markers). PANC-1 cells are a mesenchymal PDAC cell line that have demonstrated mesenchymal-to-ameboid transition under *in vitro* conditions.^34,40,66^ Plasticity in migration mode is common in PDAC and can contribute to treatment inefficacy.^34,67^ Samain et al. used a panel of pancreatic cancer cell lines across various phenotypes and genotypes to identify a mechanism for amoeboid invasion where cells move through pores and spaces within the ECM in a proteolysis-independent manner.^66^ They found high levels of CD73—an immune checkpoint—in invasive amoeboid PDAC cells that included PANC-1, and CD73 relates to RhoA-ROCK-Myosin II activation. This pathway is downstream of PI3K, which is responsive to mechanical stresses (e.g., shear, compression, tension) and important for mechanical transduction.^68^ Further study is needed to determine whether the RhoA-ROCK-Myosin II signaling pathway is activated in our model and if it is impacted by obesity signaling. However, there is a strong rationale that the amoeboid-like phenotype that we observed in PANC-1 cells is a direct result of confinement and the stiffness differences we can generate using PhotoCol®. It is likely that matrix stiffness is contributing to heterogeneous responses in our engineered tumor microenvironment by reflecting the phenotypic plasticity displayed by PDAC cells.^34,69^ Collectively, these results highlight the need to utilize model systems, like ours, that promote these phenotypes in cells to explore possibilities for new therapeutic targets.

The addition of adipocyte signaling in our system further adds to the ability to study diverse cellular responses in PDAC. Adipocytes that underwent obese stimulation in our system secreted increased IL-8 and IL-10 when compared to adipocytes differentiated in 2D to produce a lean phenotype. IL-8 can be secreted from multiple stromal populations in the tumor microenvironment. Its secretion from tumor associated macrophages has been shown to increase EMT mediated invasion in PANC-1 and BxPC-3 cells through the STAT3 pathway.^70^ IL-10 is similarly released by M2 polarized (anti-inflammatory) macrophages and increases vimentin expression in PANC-1 cells.^51^ Our results confirmed that obesity is contributing to an altered inflammatory environment in cancer, and that this inflammation is supporting mesenchymal phenotypes in PDAC cells. This relationship has been established in mouse models^21,71^, but has yet to be modelled *in vitro* with both human PDAC and adipocyte populations under fibrotic conditions.

## 4. CONCLUSIONS

Overall, this work contributes to the larger scope of understanding how obesity and fibrosis influences PDAC malignancy. Fibrotic progression of the extracellular matrix is a hallmark of PDAC and has an emerging role in inducing adipocyte dysfunction in obesity. Obesity has been well established as a risk factor for developing PDAC, but there is a need for *in vitro* model systems that focus on cell interactions that occur once the disease has initiated to answer questions about persistence and progression. We leveraged the ability to reproducibly alter biophysical properties (i.e., stiffness) of a natural hydrogel via photo-crosslinking (PhotoCol®) and provide pathological signaling from human adipocytes (i.e., obesity). Our results showed a role for obesity in modulating vimentin expression in PANC-1 cells, which relates to EMT and malignancy, and increased matrix stiffness appeared to promote an amoeboid-like phenotype as an alternative mode of invasion. Obesity signaling also enhanced vimentin expression while maintaining amoeboid morphology, which demonstrates the ability of our model to generate diverse phenotypic changes in PDAC cells *in vitro*. Overall, our study revealed an opportunity to leverage the stiffening and microstructural properties of PhotoCol® to recreate an appropriate microenvironment to study amoeboid invasion, which is understudied *in vitro*. Our work also highlights the importance of studying cell signaling dynamics under fibrotic conditions in PDAC and obesity, which is necessary for understanding fundamental interactions that drive malignancy and support the development of therapeutic interventions for PDAC for diverse patients.

## 5. EXPERIMENTAL SECTION

### 5.1 Cell culture for PANC-1 cells and adipocytes

PANC-1 cells were obtained from American Type Culture Collection (ATCC; Manassas, Virginia) were maintained according to manufacturer instructions in Dulbecco’s Modified Eagle Medium (DMEM) supplemented with 10% fetal bovine serum (Corning) and 1X penicillin-streptomycin (Gibco). Cells were used between passages 4 and 15. Mature adipocytes were obtained from differentiating human adipose-derived mesenchymal stem cells (AD-MSC; ATCC) using an Adipocyte Differentiation Toolkit (ATCC). Briefly, AD-MSCs were seeded at 18,000 cells/ cm^2^ in 6 well tissue-culture treated well plates and grown to confluence for 48 hours in mesenchymal stem cell basal medium supplemented with a penicillin-streptomycin-amphotericin B solution (ATCC). Confluent AD-MSCs were exposed to differentiation medium (Adipocyte Basal Medium with AD Supplement) for an additional 96 hours to initiate adipogenic differentiation. Cultures were then switched to adipocyte maintenance media (Adipocyte Basal Medium with ADM Supplement) for an additional 10 days to promote adipocyte maturation. Once in the maturation phase, media was changed every 3-4 days. Adipogenic differentiation was confirmed by the presence of round lipid droplets and with Oil red O staining (Millipore Sigma). Cells were used for conditioned media generation until day 30 of culture. All cells were maintained in a humidified incubator at 37°C with 5% CO_2_.

### 5.2 PANC-1 exposure to adipocyte conditioned media (indirect co-culture)

To verify that PANC-1 cells tolerated exposure to adipocyte growth medium prior to adipocyte conditioned media exposure, PANC-1 cells were exposed to varied ratios of unconditioned adipocyte growth medium to DMEM. Cells were initially plated in 2D at 10,000 cells/well in a 96-well plate and cultured in DMEM for 24 hours, after which DMEM was removed. Samples were then rinsed with 1X DPBS (no calcium, no magnesium) and overlaid with ratios of adipocyte growth medium to DMEM: 50% v/v and 75% v/v. Samples were cultured for an additional 48 hours before being evaluated with CalceinAM/Ethidium homodimer LIVE/DEAD assay (ThermoFisher) to determine percent viability. Once viability was confirmed with adipocyte growth medium, the same ratios of adipocyte conditioned medium to DMEM were added to PANC-1 cells on tissue culture plastic. Viability and cell morphology were assessed as previously described. The highest ratio of adipocyte conditioned media that supported PANC-1 viability (>80%) was used for exposure studies in 3D.

### 5.3 Preparation of PANC-1 Seeded PhotoCol® Hydrogels

Lyophilized methacrylated type I collagen (PhotoCol®, Advanced BioMatrix, Carlsbad, CA) was solubilized in sterile 20mM acetic acid to a final concentration of 8mg/mL on a rotator at 4°C. To polymerize PhotoCol® prior to photo-crosslinking, solubilized PhotoCol® was combined with neutralization solution (8% v/v; Advance BioMatrix) and lithium phenyl-2,4,6-trimethylbenzoylphosphinate (LAP) photoinitiator (2% v/v; Advanced BioMatrix). All components were thoroughly mixed on ice to prevent premature polymerization, and final concentration was 7.262 mg/mL. To fully polymerize PhotoCol® with or without cells, hydrogels were incubated at 37°C for 30 minutes. For cell seeding, sub-confluent PANC-1 cells were trypsinized (0.25%), collected, and centrifuged to form a cell pellet. After removing the supernatant and breaking up cell pellet, PANC-1 cells were resuspended in neutralized PhotoCol® (300,000 cells/mL; 80 µl sample volume) and pipetted into black walled, clear-bottom, polystyrene, 96-well plates (Corning, NY). Samples were polymerized at 37°C for 30 minutes and overlaid with PANC-1 culture media prior to incubation at 37°C with 5% CO_2_. To produce higher stiffness samples, cell-seeded PhotoCol® containing LAP photoinitiator were photo-crosslinked with 405 nm light source (3-minute exposure) 4 hours after cell seeding. Lower stiffness samples were prepared with photoinitiator but with no light exposure. For all samples, cell culture media was changed on day 1 of culture.

### 5.4 Phenotypic assessment of PANC-1 response in 2D and 3D

Samples were fixed with 4% paraformaldehyde (ThermoFisher Scientific, Waltham, MA) and permeabilized with 0.1% Triton X-100 (Sigma Aldrich). To evaluate morphological changes, samples were stained with Alexa Fluor 488 Phalloidin (ThermoFisher Scientific) to visualize F-actin. For immunostaining to evaluate indicators of epithelial-to-mesenchymal transition (EMT) and invasiveness, fixed samples were blocked with 5% bovine serum albumin (BSA) in 1X PBS and incubated with mouse monoclonal anti-vimentin primary (Millipore Sigma, 1:1000 in 1% BSA) overnight at 4°C. Samples were then rinsed with 1X PBS and incubated with Alexa Fluor-conjugated secondary antibodies: 488 Phalloidin and 594 Goat anti-Mouse IgG (H+L) Highly Cross-Adsorbed Secondary Antibody (ThermoFisher Scientific, 1:1000). All samples were counterstained with 4′,6-diamidino-2-phenylindole (DAPI; ThermoFisher Scientific) to visualize cell nuclei.

To qualitatively and quantitatively assess phenotypic changes, stained samples were imaged with a Keyence BZX810 All-in-One Fluorescence Microscope using a Plan Fluorite 20x LD objective (KEYENCE Corp. of America, Itasca, IL). Maximum projection images were used for analysis (100-200 µm depth, 7 µm pitch). Quantitative image analysis was performed using CellProfiler™ (Broad Institute, Cambridge, MA) to assess change in cell shape metrics associated with cell elongation (extent, eccentricity) and cell spread (compactness, solidity, form factor) based off work from Baskaran et al.^33^ Extent is the area of an object divided by the area of a bounding box that encompasses the entire object. Eccentricity is a measure of how much an object deviates from being a perfect circle. Compactness is the mean squared distance of an object’s pixels from the centroid divided by the area. A value of 1 is representative of a perfect circle, while values greater than one indicate irregular shape boundaries or objects with holes. Solidity is the proportion of the object area to convex hull area. Form factor is equal to 4*π*Area/Perimeter^2^. Relative vimentin expression was expressed as integrated density of vimentin staining in identified cells.

### 5.5 Preparation and verification of adipocytes with obese phenotype

Adipocyte differentiation on 2D surfaces was representative of mature healthy adipocytes that make up lean fat. To generate adipocytes that were more representative of an obese phenotype, adipose-derived MSCs were differentiated within a 6 mg/mL type I collagen hydrogel (TeloCol, Advanced BioMatrix) to recapitulate properties of obese tissue. Collagen samples with encapsulated MSCs were plated in 96 well plates and allowed to polymerize for 30 min at 37°C, after which they were incubated in regular MSC culture medium for 48 hours before initiating differentiation with the Adipocyte Differentiation Toolkit (ATCC). Samples were then switched to induction medium for 96 hours, followed by maintenance medium for the remainder of differentiation. Encapsulated adipocytes grew for 14 days to full maturity, and their medium was collected for secretome characterization and use in experiments until day 25 of culture. Conditioned media was collected from mature adipocytes (> 14 days of differentiation or visible lipid droplet formation) every 3 days and centrifuged (300 x g) for 5 minutes to remove cell debris. Conditioned media then sterile filtered and stored at -80°C for use in subsequent experiments. Obese phenotype was determined using a Human Obesity Array C1 (RayBiotech, Peachtree Corners, GA) to detect the presence of key adipokines. Assay membranes were blocked for 30 minutes at room temperature and conditioned media samples from lean and obese adipocyte cultures were incubated with membranes overnight at 4°C. Post incubation, biotinylated antibodies were added to the membranes followed by horseradish peroxidase (HRP)-Streptavidin prior to chemiluminescence detection. Imaging was done with an Azure 600 Gel Doc (Azure Biosystems, Dublin, CA). Data extraction and densiometry analysis was performed in ImageJ using the Protein Array Analyzer Plugin (Gilles Carpentier). Blots were normalized using positive control/reference and negative control spots on each blot using the manufacturer supplied analysis tool (RayBiotech).

### 5.6 Statistical Analysis

Statistical analysis was performed with GraphPad Prism. For 2D cell morphology, a one-way ANOVA with Dunnett’s multiple comparisons was performed to compare the mean of each group to the mean of the control group (DMEM). For 3D morphology and vimentin expression, samples within one stiffness level were compared with a one-way non-parametric ANOVA (Kruskal-Wallis) with Dunn’s multiple comparisons. To compare shape metrics and vimentin expression between stiffness levels for one media formulation, a Mann-Whitney U-test was used. Averages of dot blot densitometry measurements were compared with an un-paired t-test. In all cases, results were considered significantly different when p < 0.05.

## ACKNOWLEDGMENTS

The authors acknowledge Hannah Borges, Madison Sanborn, and Eleanor Finberg for their assistance with cell culture.

## AUTHOR CONTRIBUTIONS

Author contributions: Conceptualization and designed experiments, A.E.J. and C.F.W.; Methodology, A.E.J., J.F.N., T.L.F., B.N.K.R.; Formal analysis, A.E.J. and J.F.N.; Figure preparation, A.E.J. and C.F.W.; Resources, C.F.W.; Original draft preparation, A.E.J. and C.F.W.; Review and editing, A.E.J. and C.F.W. Authors have read and agreed to the published version of the manuscript.

